# Kinetic scheme of myosin phosphorylation by ZIP kinase

**DOI:** 10.1101/2025.08.15.670483

**Authors:** Mayu Yamaguchi, Reiko Nakagawa, Linh T. Tran, Yoshihiro Shimizu, Makito Miyazaki

**Affiliations:** Faculty of Agriculture, Kyoto University, Kitashirakawa Oiwake-cho, Sakyo-ku, Kyoto 606-8502, Japan; RIKEN Center for Biosystems Dynamics Research, 2-2-3 Minatojima-minamimachi, Chuo-ku, Kobe, Hyogo 650-0047, Japan; RIKEN Center for Integrated Medical Sciences, 1-7-22 Suehiro-cho, Tsurumi-ku, Yokohama, Kanagawa 230-0045, Japan; Graduate School of Frontier Biosciences, Osaka University, 1-3 Yamadaoka, Suita, Osaka 565-0871, Japan; Graduate School of Medicine, Science and Technology, Shinshu University, 3-1-1 Asahi, Matsumoto, Nagano 390-8621, Japan; PRESTO, JST, 4-1-8 Honcho, Kawaguchi, Saitama 332-0012, Japan; The Hakubi Center for Advanced Research, Kyoto University, Yoshida Honmachi, Sakyo-ku, Kyoto 606-8501, Japan; Department of Physics, Graduate School of Science, Kyoto University, Kitashirakawa Oiwake-cho, Sakyo-ku, Kyoto 606-8502, Japan

**Author notes:** These authors are equally contributed to this work (Mayu Yamaguchi, Reiko Nakagawa).

**Keywords:** Myosin, Phosphorylation, Contractility, Protein kinase

## Abstract

Zipper-interacting protein kinase (ZIPK) is a ubiquitous serine/threonine protein kinase that plays pivotal roles in regulating cell motility, division, and smooth muscle contractility through phosphorylation of myosin. In this study, we systematically investigated the phosphorylation reactions of smooth muscle myosin (SMM) by ZIPK. We found that ZIPK phosphorylates MRLC sequentially, first at Ser^19^ and then at Thr^18^, determined by quantitative mass spectrometry analysis on wild-type MRLC. Analysis on phosphomimic and unphosphorylatable MRLC mutants indicates that the phosphorylation rate at Ser^19^ on unphosphorylated MRLC is 1.5 times faster than that at Thr^18^ on Ser^19^-phosphorylated MRLC. Comparison between SMM and isolated MRLC revealed that the phosphorylation rate of SMM is slower than that of isolated MRLC. To dissect the molecular mechanism responsible for this difference, we measured interactions between ZIPK and SMM by co-sedimentation assay. The result suggests that the C-terminal domain of ZIPK interacts with the heavy chain of SMM, and as a result, competitive binding of ZIPK to MRLC and the myosin heavy chain suppresses phosphorylation of SMM compared to isolated MRLC. By incorporating the kinetic and dissociation constants obtained from mutant analysis and co-sedimentation assays, respectively, a simple kinetic model accurately reproduced the time courses of phosphorylation for both isolated MRLC and SMM. This provides systemic insight into the regulatory mechanism of myosin contractility by ZIPK.

## Introduction

Myosin II is the primary force generator of the actin cytoskeleton, which regulates various cellular functions, including cell motility, division, and muscle contractility[1, 2]. Myosin molecules oligomerize to form myosin filaments, which markedly increases the binding lifetime on actin filaments, and enables to slide and generate tensile force on the filaments coupled with ATP hydrolysis[1]. Activation of smooth muscle myosin (SMM) and non-muscle myosin (NMM) requires phosphorylation of the myosin regulatory light chain (MRLC, also called LC20)[2, 3]. It has been determined that MRLC has two phosphorylation sites that contribute to the activation of myosin[4]. Monophosphorylation of MRLC at serine^19^ (Ser^19^) is sufficient to stimulate the assembly of myosin filaments[5], actin-activated Mg^2+^-ATPase[6], and force generation[7]. Diphosphorylation of MRLC at threonine^18^ (Thr^18^) and Ser^19^ further enhances myosin filament formation[5, 8] and actin-activated Mg^2+^-ATPase[6, 9], however, it does not increase the gliding velocity of actin filaments when compared to monophosphorylated myosin[9].

Several protein kinases have been identified that phosphorylate MRLC at Thr^18^ and Ser^19^. Myosin light chain kinase (MLCK) is a ubiquitous protein kinase found in smooth muscle and non-muscle cells. This kinase is involved in the regulation of various cellular processes, including cell spreading, migration, cytokinesis, and the regulation of smooth muscle contractility through MRLC phosphorylation[3, 10]. MLCK predominantly phosphorylates MRLC at Ser^19^, with phosphorylation at Thr^18^ occurring at a much slower rate[4]. Activation of its kinase activity requires the binding of Ca^2+^/calmodulin[11, 12]. Rho-associated coiled-coil kinase (ROCK) is one of the other ubiquitous serine/threonine protein kinases implicated in a wide range of cellular functions through the phosphorylation of various substrates, including MRLC. Similar to MLCK, ROCK primarily phosphorylates MRLC at Ser^19^[13], with its kinase activity regulated by the binding of GTP-bound active Rho[14].

In addition to MLCK and ROCK, the zipper-interacting protein kinase (ZIPK, also called DAPK3), initially identified as a serine/threonine kinase that mediates apoptosis[15], also phosphorylates MRLC[16]. Growing evidence suggests that ZIPK plays an important role in the regulation of cell motility[17], cytokinesis[18, 19], reorganization of the actin cytoskeleton in non-muscle cells[20], and smooth muscle contractility[21, 22] through phosphorylation of MRLC. ZIPK is composed of a protein kinase domain in its N-terminus and a leucine zipper motif in its C-terminus[15] (**Fig. 1**). ZIPK is a member of a large family of death-associated protein kinases (DAPK) and shares 83% identity at the amino acid level with the kinase domain of DAPK. However, it shows no homology in its C-terminal domain to the other members of the DAPK family[23, 24]. Unlike major DAPK family members, ZIPK does not contain a death domain or a calmodulin-binding domain, and its kinase activity is regulated independently of Ca^2+^[21, 23, 24].

**Figure 1.**
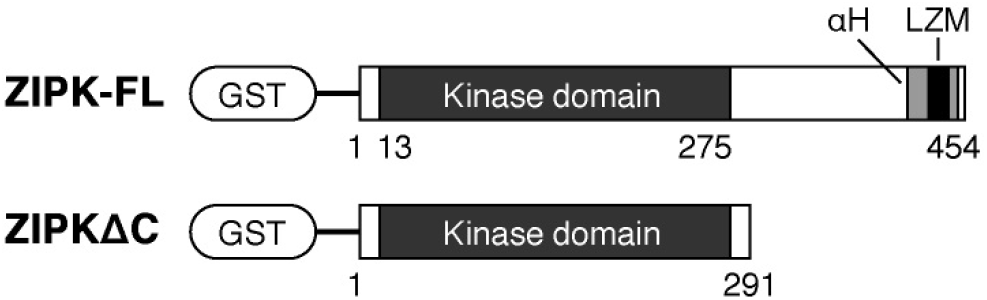
Structures of the recombinant ZIPK used in this study. Full-length human ZIPK (ZIPK-FL) and the C-terminal deletion mutant (ZIPKΔC) were fused to a glutathione S-transferase (GST)-tag. The N-terminus of ZIPK contains a serine/threonine protein kinase domain (13–275 a.a.)[15]. The C-terminus is predicted to form an *α*-helical structure (*α*H; 413–450 a.a.) containing a leucine zipper motif (LZM; 427–441 a.a.)[15].

ZIPK phosphorylates MRLC at both Ser^19^ and Thr^18^, confirmed firstly by phospho-amino acid analysis[16], followed by mutational analysis of MRLC[21]. Unlike MLCK and ROCK, ZIPK has been proposed to phosphorylate Ser^19^ and Thr^18^ with nearly identical rates in a random order, based on the good agreement between experimental time courses of wild-type MRLC phosphorylation and predictions from a random phosphorylation model[21]. In contrast, the original report of ZIPK suggested a sequential mechanism, i.e., phosphorylation first at Ser^19^ and then at Thr^18^, based on two-dimensional phosphopeptide mapping and comparisons with the pattern of MLCK-phosphorylated MRLC[16]. Therefore, the reaction scheme remains under debate, and further investigation is required to determine the kinetic parameters and clarify the phosphorylation mechanism. Moreover, it has been known that the phosphorylation rate of SMM is slower than that of isolated MRLC[21, 25], yet the molecular mechanism underlying this difference remains unresolved.

Here, we report that ZIPK phosphorylates MRLC sequentially, first at Ser^19^ and then at Thr^18^, as determined by quantitative mass spectrometry analysis on wild-type MRLC. Gel electrophoresis of phosphomimic and unphosphorylatable MRLC mutants indicates that the phosphorylation rate at Ser^19^ on unphosphorylated MRLC is 1.5 times faster than that at Thr^18^ on Ser^19^-phosphorylated MRLC. Co-sedimentation assay of ZIPK and SMM suggests that the C-terminus of ZIPK interacts with the heavy chain of SMM. Based on this observation, we propose that competition between the binding of the C-terminal domain to the myosin heavy chain and the binding of the N-terminal kinase domain to MRLC suppresses the phosphorylation of SMM, when compared to isolated MRLC. Using the Michaelis constants and the catalytic rate constants of MRLC phosphorylation by ZIPK, and the dissociation constants between SMM and ZIPK determined from independent experiments, the time courses of the phosphorylation reactions of isolated MRLC and SMM were reproduced by a simple kinetic model with high accuracy.

## Materials and Methods

### Buffer

All experiments were performed using A50 buffer (50 mM HEPES-KOH pH 7.6, 50 mM KCl, 5 mM MgCl2, 1 mM EGTA)[33], unless otherwise stated.

### Protein preparation

Recombinant human ZIPK was prepared according to our previous report[28]. Briefly, full-length ZIPK (ZIPK-FL; 1–454 a.a.) and the C-terminal deletion mutant (ZIPKΔC; 1–291 a.a.) were cloned into pGEX-5X and expressed in *E. coli* (Rosetta 2 (DE3), Merck Millipore) at 25°C for 1 hour in the presence of 1 mM IPTG. The GST-tagged proteins were purified using a glutathione Sepharose column (GSTrap HP, GE Healthcare), followed by overnight dialysis against A50 buffer containing 1 mM DTT at 4°C. Samples were centrifuged at 627, 000 × *g* for 20 min at 4°C to remove aggregates, snap-frozen in liquid nitrogen, and stored at −80°C.

Recombinant human myosin regulatory light chains isoform 2 (MRLCs) with various mutations (wild-type, T18A, T18D, S19A, S19D, T18DS19D) were cloned into pGEX-6P, expressed in *E. coli*. (Rosetta 2 (DE3), Merck Millipore) at 16°C for 21–22 hours in the presence of 1 mM IPTG. The GST-tagged proteins were purified using a glutathione Sepharose column (GSTrap HP, GE Healthcare) and digested by PreScission Protease (GE Healthcare) overnight at 4°C. The digested samples were further purified using a size exclusion column (Superdex 200 Increase 10/300 GL, GE Healthcare) with A50 buffer containing 1 mM DTT, snap-frozen in liquid nitrogen, and stored at −80°C.

Unphosphorylated smooth muscle myosin (SMM) was purified from chicken gizzards as previously reported[29]. The extracted sample was further dialyzed at 4°C overnight, against A50 buffer containing 1 mM DTT and 0.5 mM ATP. After the addition of 0.5 mM fresh ATP to the dialyzed sample, it was centrifuged at 20, 000 × *g* for 10 min at 4°C to remove aggregates and contaminating phosphorylated myosin. The supernatant was snap-frozen in liquid nitrogen and stored at −80°C. To isolate the myosin regulatory light chain (MRLC, or LC20) and myosin essential right chain (MELC, or LC17) from SMM, the supernatant was mixed with an identical volume of 8 M urea dissolved in 50 mM Tris-HCl pH 8.0, 550 mM KCl, 10 mM EDTA, and 5 mM DTT, and incubated for 15 min at room temperature. After incubation, the sample was diluted with 10 volumes of cold distilled water, and centrifuged at 2, 000 × *g* for 30 min at 2°C to precipitate the myosin heavy chain. The supernatant containing equimolar amounts of MRLC and MELC was dialyzed against A50 buffer containing 1 mM DTT at 4°C, snap-frozen in liquid nitrogen, and stored at −80°C.

The purity of proteins was confirmed by SDS-PAGE. The protein concentrations were determined using a Protein Assay Kit (500-0006, Bio-Rad), and molecular weights of 474,000 Da for SMM (hexamer: 2 myosin heavy chains (MHC) + 2 myosin regulatory light chains (MRLC) + 2 myosin essential light chains (MELC)), 37,000 Da for the mixture of MRLC and MELC, or the molecular weights estimated using Benchling (Benchling Inc.).

### Phosphorylation rate analysis

The phosphorylation rates of MRLC and SMM by ZIPK were measured using urea/glycerol PAGE[26]. The reaction mixture (0.5–16 *μ*M MRLC or 1 *μ*M SMM, 1–100 nM ZIPK, 0.05 mg ml/BSA, 1 mM ATP, 10 mM phosphocreatine, 0.1 mg/ml creatine phosphokinase, 10 mM DTT in A50 buffer) was incubated at 25 ± 1°C using a water bath (AB-1600, ATTO) or a heat block (ThermoMixer, Eppendorf) for 0–90 min, and the reaction terminated by the addition of 1.1 g/ml urea. Unphosphorylated (0P), monophosphorylated (1P), and diphosphorylated (2P) MRLCs were separated by urea/glycerol PAGE (10 mA, 230 min). The gel was stained using the SYPRO Ruby protein gel stain (S12000, Invitrogen). The gel images were captured by Gel Doc EZ Imager (Bio-Rad) and the band intensities were analyzed using Image Lab (Bio-Rad).

### Synthetic Peptide Standards

Synthetic Ser^19^ or Thr^18^ mono-phosphorylated peptides were purchased from Eurofins Genetics. The peptides were desalted using a MonoSpin C18 column (GL Science) following the manufacturer’s protocol. After desalting, the peptides were dried and dissolved in a solution composed 2% acetonitrile and 0.1% trifluoroacetic acid. The peptide solutions were adjusted to a final concentration of 10 fmol/*μ*L for standard curve measurements. For linear regression analysis, varying amounts of synthetic phosphorylated peptides (0, 1, 2, 5, 10, 20, and 50 fmol) were injected sequentially. The quantification was performed using the extracted ion chromatogram (XIC) of the MS2 fragment ion at 579.3 *m/z*, which corresponds to the most intense double-positive ion of neutral loss (−98) derived from the monophosphorylated peptide precursor ion at 628.3 *m/z*.

### LC-MS/MS spectrometry analysis

The phosphorylation ratio of wild-type MRLC at Ser^19^ and Thr^18^ was measured by mass spectrometry analysis. The reaction mixture (2 *μ*M MRLC, 5 nM ZIPK, 0.05 mg/ml BSA, 1 mM ATP, 10 mM phosphocreatine, 0.1 mg/ml creatine phosphokinase, 10 mM DTT in A50 buffer) was incubated for 10 min at 25 ± 1°C using a water bath (AB-1600, ATTO). After the incubation, the reaction was immediately stopped by adding 2% (w/v) of sodium dodecyl sulfate (SDS) and 5% (v/v) of 2-mercaptoethanol, followed by incubation at 98°C for a few minutes. The phosphorylated proteins were reduced with 10 mM Tris(2-carboxyethyl) phosphine hydrochloride (TCEP) and alkylated by incubation with 55 mM iodoacetamide (IAA) for 30 min at room temperature in the dark. The protein solutions were then digested with Lys-C using the SP3 method[30], and the resulting peptide solutions were collected and dried using a SpeedVac. Additional digestion with pepsin was subsequently performed. The Lys-C and pepsin double digests were desalted using in-house made C18 Stage-Tips.

Mass spectra were acquired with an Orbitrap Eclipse mass spectrometer (Thermo Fisher Scientific) coupled to a nanoflow UHPLC system (Vanquish; Thermo Fisher Scientific). The peptides were loaded onto a C18 trap column (PepMap Neo Trap Cartridge, ID 0.3 mm × 5 mm, particle size 5 *μ*m, Thermo Fisher Scientific) and fractionated through the C18 analytical column (Aurora, ID 0.075 mm × 250 mm, particle size 1.7 *μ*m, Ion Opticks). The peptides were eluted at a flow rate of 300 nL/min using the following gradient: 0% to 2% solvent B over 1 min, 2% to 5% over 2 min, 5% to 16% over 19.5 min, 16% to 25% over 10 min, 25% to 35% over 4.5 min, a sharp increase to 95% over 4 min, held at 95% for 5 min, and finally re-equilibrated at 5%. Solvent A and B compositions were 100% H2O with 0.1% formic acid and 100% acetonitrile with 0.1% formic acid, respectively.

For the determination of phosphorylation sites, the Orbitrap Eclipse was operated in data-dependent acquisition (DDA) mode with a 3-second cycle time, spray voltage of +2000 V, capillary temperature of 275°C, funnel RF lens at 30%, and lock masses set at 445.12003, 519.13882, and 536.16537. The MS1 scan was collected at a resolution of 120,000 at 200 *m/z*, over a mass range of 375–1500 *m/z*, using a standard AGC and a maximum injection time of 50 ms. MS/MS was triggered from precursors with intensity above 20,000 and charge states of 2–7. Quadrupole isolation width was set to 1.6 *m/z*, with normalized HCD energy of 30%, and resulting fragment ions recorded in the Orbitrap. MS2 scans were collected at a resolution of 30,000 at 200 *m/z*, with a standard AGC target and a maximum injection time of 54 ms. Dynamic exclusion was set to 20 seconds.

For parallel reaction monitoring (PRM) analysis, the Orbitrap Eclipse was operated in data-independent acquisition PRM mode, spray voltage of +2000 V, capillary temperature of 275°C, and lock masses set at 445.12003, 519.13882, and 536.16537. A survey full-scan MS (from *m/z* 375–1500) was acquired in the Orbitrap at a resolution of 120,000 at 200 *m/z*. A targeted list of precursor ions was set at 628.3 *m/z*, isolated and fragmented using HCD with normalized collision energy of 30%, and detected at a mass resolution of 30,000 at 200 *m/z*. The data were subsequently analyzed using Skyline to obtain the total area for absolute quantification[31].

The DDA raw data files were searched against the amino acid sequence of hMRLC with the Common Repository of Adventitious Proteins (cRAP, ftp://ftp.thegpm.org/fasta/cRAP) using Proteome Discoverer 2.5 software (Thermo Fisher Scientific) with the MASCOT ver. 2.8 search engine, with a false discovery rate (FDR) set at 0.01. MS and MS/MS mass tolerances were set to 10 ppm and 0.02 Da, respectively. Cleavage sites were set at the C-termini of lysine, phenylalanine, leucine, tryptophan, and tyrosine, which occur as a result of double digestion with Lys-C and pepsin, and a maximum of two cleavage misses were allowed. Carbamidomethylation of cysteine was set as a fixed modification, while oxidation of methionine, phosphorylation of serine, threonine, and tyrosine, and acetylation of protein N-termini were set as variable modifications.

### Co-sedimentation assay

To measure the binding affinity of ZIPK for SMM, a co-sedimentation assay was performed. The reaction mixture (1 *μ*M smooth muscle myosin, 125–500 nM ZIPK, 0.05 mg/ml BSA, 1 mM ATP, 10 mM phosphocreatine, 0.1 mg/ml creatine phosphokinase, 10 mM DTT in A50 buffer) was incubated at 25 ± 1°C using a water bath (AB-1600, ATTO) for 60 min. A small aliquot of the reaction mixture was taken, and the rest was centrifuged at 627, 000 × *g* for 20 min at 25°C. Identical aliquots of the uncentrifuged whole sample (W) and the supernatant (S) were subjected to SDS-PAGE (5–20% gradient precast gel, ATTO). The gel was stained using SYPRO Ruby protein gel stain (S12000, Invitrogen). The gel images were captured by Gel Doc EZ Imager (Bio-Rad) and the band intensities were analyzed using Image Lab (Bio-Rad).

## Results

### Determination of the phosphorylation sites of MRLC by ZIPK

We first analyzed the phosphorylation time course of wild-type human MRLC by ZIPK-FL. Two micromolar of the wild-type MRLC were incubated with 1, 5, 20 nM of the full-length human ZIPK (ZIPK-FL) for various time periods at 25°C, and the reactions were terminated by the addition of urea. The samples were then subjected to urea/glycerol poly-acrylamide gel electrophoresis (urea/glycerol PAGE) to separate unphosphorylated and phosphorylated MRLCs[26, 27]. The urea/glycerol PAGE clearly separated unphosphorylated (0P), monophosphorylated (1P), and diphosphorylated (2P) MRLCs (**Fig. 2A**). The phosphorylation rate increased in a dose-dependent manner with the ZIPK-FL concentration (**Fig. 2B**).

**Figure 2.**
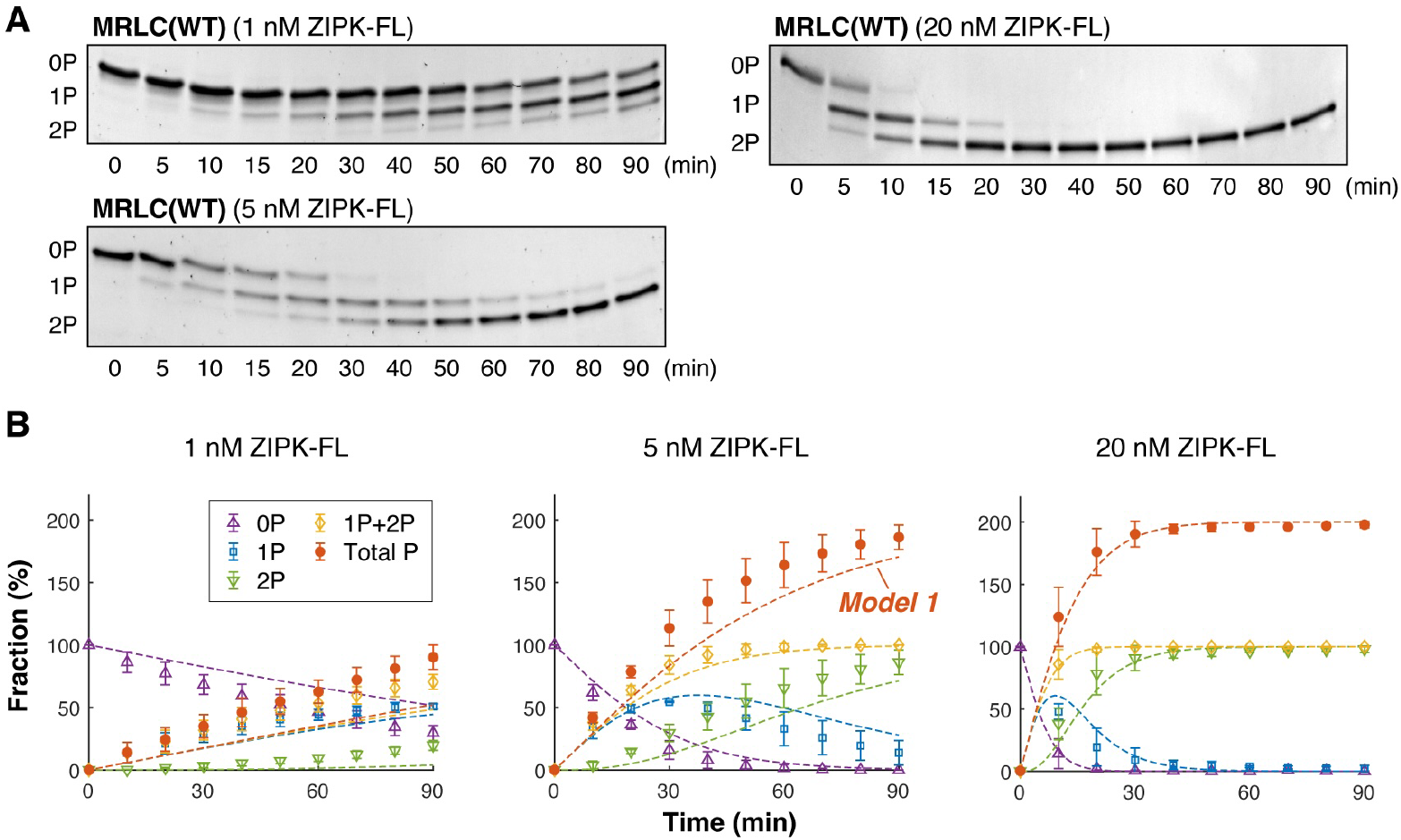
Phosphorylation reactions of wild-type MRLC by ZIPK-FL quantified by urea/glycerol PAGE. **A**. Images of urea/glycerol PAGE. Two micromolar of wild-type human MRLC was incubated with 1, 5, or 20 nM of ZIPK-FL in the presence of 1 mM ATP for the indicated times at 25°C. Unphosphorylated (0P), monophosphorylated (1P), and diphosphorylated (2P) MRLCs were separated by gel electrophoresis. **B**. Time course of the phosphorylation reactions. Fractions of 0P, 1P, 2P, and 1P+2P MRLCs, determined from densitometry of the urea/glycerol PAGE shown in A, were plotted. Total P indicates the total phosphorylation level, i.e., total P (%) = 1P (%) + 2 × 2P (%). Three independent experiments were performed. The plots and error bars indicate the mean values and SDs, respectively. The dashed lines represent the time courses predicted from **Model 1** (Eqs. (1) and (2), **Fig. 5B**, right) using the kinetic constants summarized in **Table 1**.

To determine the phosphorylation sites of MRLC by ZIPK, we next prepared human MRLC mutants in which Ser^19^ (S19) or Thr^18^ (T18) were mutated to alanine (A) or aspartic acid (D) to mimic unphosphorylated and phosphorylated states, respectively. Two micromolar of the respective mutants were incubated with 5 nM of the full-length human ZIPK (ZIPK-FL) for various time periods at 25°C, and the reactions were terminated by the addition of urea. The samples were then subjected to urea/glycerol PAGE. All four mutants had only two states (**Fig. 3A**, S19A, S19D, T18A, and T18D), suggesting that only Ser^19^ and Thr^18^ are the phosphorylation sites, and that the remaining amino acids are not phosphorylated by ZIPK. This was confirmed using an MRLC mutant in which both Ser^19^ and Thr^18^ were mutated to aspartic acid to mimic the diphosphorylated state. This mutant did not show a band shift by ZIPK-FL treatment (**Fig. 3A**, T18DS19D), verifying that only Ser^19^ and Thr^18^ are phosphorylated by ZIPK. These results are consistent with a previous study conducted using an autoradiographic method with [*γ*-^32^P]ATP[21].

**Table 1.** The kinetic constants for the phosphorylation of Thr^18^ and Ser^19^. The values were obtained by nonlinear least-square fitting of the Michaelis–Menten equation to the raw data plots (**Fig. 5A** and **Fig. 8**).

|  | ZIPK-FL |  | ZIPK-ΔC |  |
| --- | --- | --- | --- | --- |
|  | Thr<sup>18</sup> | Ser<sup>19</sup> | Thr<sup>18</sup> | Ser<sup>19</sup> |
| $K_m$ (μM) | 4.2 | 4.2 | 2.5 | 2.5 |
| $k_{cat}$ (sec<sup>-1</sup>) | 0.40 | 0.60 | 0.63 | 0.95 |

**Figure 3.**
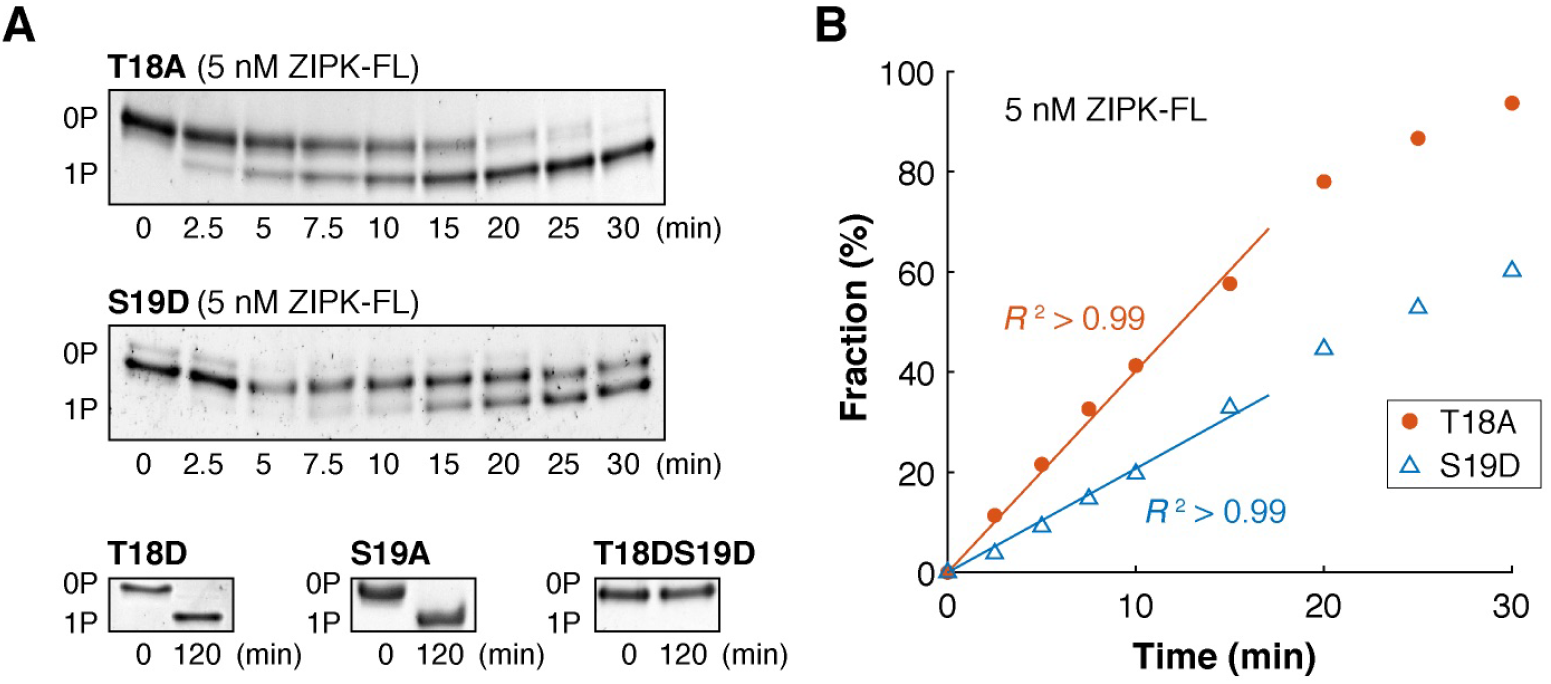
Determination of the phosphorylation sites of MRLC by ZIPK. **A**. Images of urea/glycerol PAGE. Two micromolar of unphosphorylatable or phosphomimic mutants of human MRLC were incubated with 5 nM ZIPK-FL in the presence of 1 mM ATP for the indicated times at 25°C. Unphosphorylated (0P) and monophosphorylated (1P) MRLCs were separated by the gel electrophoresis. **B**. Analysis results of the urea/glycerol PAGE shown in A (T18A and S19D). The intensities of the 0P and 1P bands were analyzed, and the fraction of 1P bands were plotted against the reaction time. The solid lines indicate the least-squares fitting of a linear function for the experimental data between 0 and 15 min. The lines mostly overlap with the experimental data (*R*^2^ > 0.99), showing that the phosphorylation reaction can be approximated as a linearly increasing phase in this period.

### Determination of the phosphorylation reaction scheme of MRLC by ZIPK through quantitative mass spectrometry analysis

We next performed quantitative mass spectrometry analysis of wild-type MRLC. Two micromolar of wild-type MRLC was treated with 5 nM ZIPK-FL for 10 min at 25°C, then the phosphorylation reaction was immediately terminated by adding SDS. This sample was subjected to the mass spectrometry analysis. It was estimated from the urea/glycerol-PAGE analysis that 34.3 ± 9.3% (*n* = 3) of MRLC was monophosphorylated (1P) and 3.7 ± 3.2% *n* = 3) of MRLC was diphosphorylated (2P) in this condition (**Fig. 2B**). Therefore, the majority of phosphorylated proteins must be in the 1P state.

Identification of monophosphorylation sites and quantification of phosphorylation levels were performed by mass spectrometric analysis of peptides generated through double-digestion with pepsin and LysC. We focused on the peptide with the sequence RPQRATSNVF and two synthetic monophosphorylated peptides (RPQRApTSNVF and RPQRATpSNVF, 628.3 *m/z*) were prepared. Comparison of their retention times and MS/MS analyses confirmed that phosphorylation occurs at Thr^18^ and Ser^19^ (**Fig. 4, Supplementary Figure 1**). Quantification was independently performed twice using calibration curves prepared from synthetic monophosphorylated peptides. As a result, the ratio of monophosphorylation levels at Thr^18^ to Ser^19^ was calculated as 1:35.7 and 1:34.2 in two biological replicates, respectively, indicating that the phosphorylation level at Ser^19^ is approximately 35 times higher than that at Thr^18^. This quantitative analysis result provides direct evidence that ZIPK-FL predominantly phosphorylates first at Ser^19^ and then at Thr^18^ in a sequential manner.

**Figure 4.**
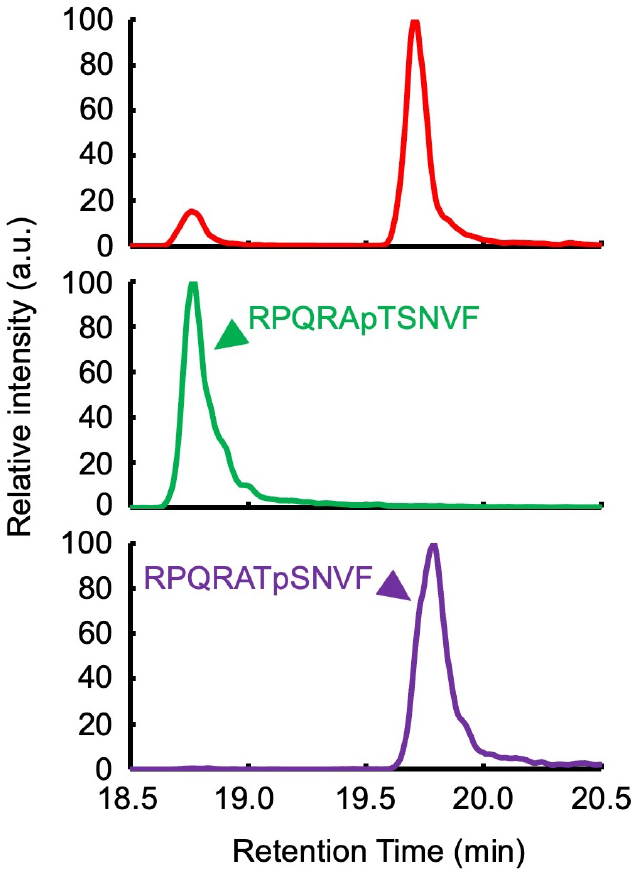
Identification of monophosphorylation sites and quantification of phosphorylation levels of wild-type MRLC by ZIPK-FL using mass spectrometry. **A**. The monophosphorylated peptides (RPQRApTSNVF and RPQRATpSNVF, *m/z* 628.3), a double digest of wild-type human MRLC with Lys-C and pepsin, were analyzed by mass spectrometry. Top: Ion chromatogram of MRLC mono-phosphorylated peptide 10 min after the phosphorylation reaction. Two peaks were observed at 18.8 min and 19.7 min. Middle: Ion chromatogram of the synthetic T18-mono-phosphorylated peptide, with a peak detected at 18.8 min. Bottom: Ion chromatogram of the synthetic S19-mono-phosphorylated peptide, with a peak detected at 19.7 min.

### Determination of the kinetic constants for the phosphorylation reactions of MRLC by ZIPK

To determine the kinetic constants of the phosphorylation reactions of MRLC by ZIPK, we used MRLC mutants in which Thr^18^ (T18) or Ser^19^ (S19) were mutated to alanine (A) or aspartic acid (D) to mimic unphosphorylated MRLC (0P-MRLC) and monophosphorylated MRLC at Ser^19^ (1P-MRLC), respectively. The extent of phosphorylation at Ser^19^ for the T18A mutant and at Thr^18^ for the S19D mutant was determined by band densitometry analysis, whereby the fraction of phosphorylated MRLC mutant was calculated by dividing the intensity of the monophosphorylated band by the sum of intensities of unphosphorylated and monophosphorylated bands. The fraction of phosphorylated MRLC increased linearly for the first 15 min in the presence of 5 nM ZIPK-FL for the two mutants (**Fig. 3B**). Therefore, we fixed the ZIPK-FL concentration and reaction time at 5 nM and 5 min, respectively, and the initial phosphorylation rates of the two mutants were measured at various MRLC concentrations, ranging from 1 *μ*M to 16 *μ*M (**Fig. 5A**). We found that the phosphorylation rate of T18A was slightly higher than that of S19D, indicating that the phosphorylation rate of 0P-MRLC at Ser^19^ was slightly higher than that of 1P-MRLC at Thr^18^ in the wild-type MRLC.

**Figure 5.**
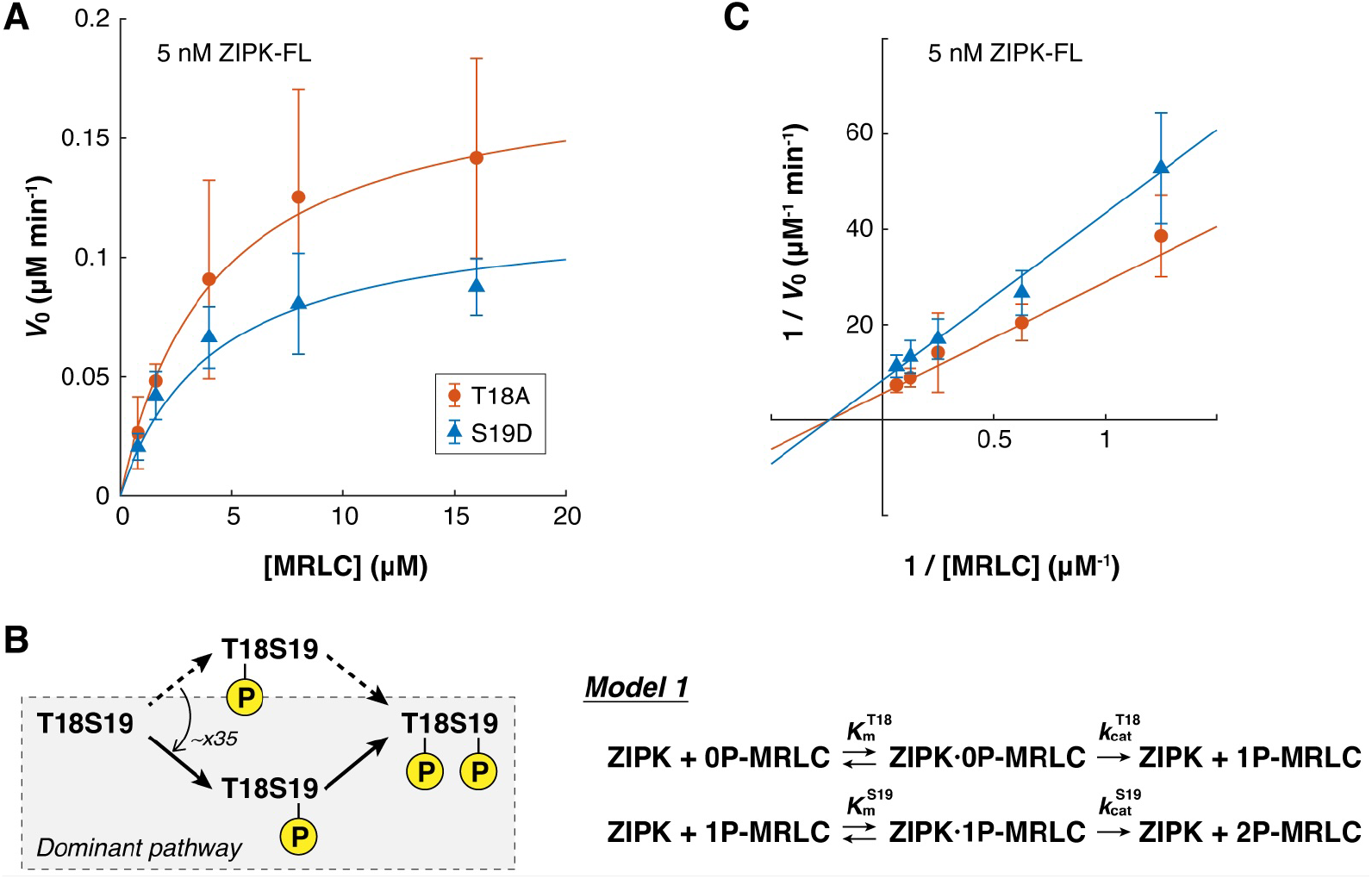
Determination of the kinetic constants for the phosphorylation reactions of MRLC by ZIPK-FL. **A**. Phosphorylation rates *V*0 of four MRLC mutants at various MRLC concentrations in the presence of 5 nM ZIPK-FL and 1 mM ATP. The mixtures were incubated at 25°C for 5 min, and the reaction rates were estimated from densitometry of the urea/glycerol PAGE (see **Fig. 3**). The mean values of the independent experiments were plotted (*n* = 3– 5). Error bars indicate the standard deviations (SDs). The solid lines indicate the nonlinear fitting of Michaelis–Menten reaction model (Eqs. (1) and (2)) using 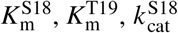, and 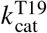 as the fitting parameters. The kinetic constants determined through the nonlinear fittings are summarized in **Table 1. B**. Reaction pathways (left), and the model (right). **C**. Lineweaver–Burk plots of A. Mean values are plotted (*n* = 3–5). Error bars indicate SDs. The lines were drawn using the kinetic constants determined from the fitting in A.

Based on the results of these mutant analyses and the mass spectrometry analysis on the wild-type MRLC, we propose that the phosphorylation reactions of MRLC by ZIPK can be approximated by two sequential Michaelis–Menten reactions (**Model 1**; **Fig. 5B**, right):

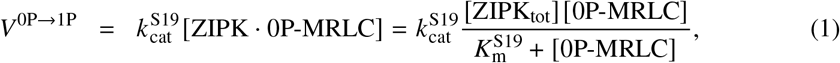

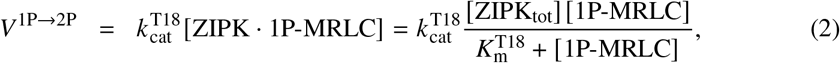

where [0P-MRLC] and [1P-MRLC] indicate the concentrations of unphosphorylated MRLC (0P-MRLC) and monophosphorylated MRLC at Ser^19^ (1P-MRLC), respectively. The catalytic rate constants *k*_cat_ and Michaelis constants *K*_m_ in Eqs. (1) and (2) were determined from the nonlinear least-square fitting of the Michaelis–Menten equation to the plots (**Fig. 5A**), as summarized in **Table 1**. The obtained *K*_m_ values are identical between T18A and S19D (4.2 *μ*M). This result suggests that the affinities of ZIPK-FL for 0P-MRLC and 1P-MRLC are comparable, and the observed difference in the reaction rates *V*_0_ is primarily atrributable to the difference in the catalytic rate constants *k*_cat_ of Ser^19^ and Thr^18^ (0.60 s^−1^ and 0.40 s^−1^, respectively). We also confirmed that the lines drawn using the obtained values were in good agreement with the Lineweaver–Burk plots (**Fig. 5C**).

### Comparison of the phosphorylation reactions between free MRLC and MRLC bound on the myosin heavy chain

To test the accuracy of **Model 1** (**Fig. 5B**, right), the time course of the phosphorylation reactions was simulated using Eqs. (1) and (2), and the kinetic parameters determined from the mutant experiments (**Table 1**). This model predicted the experimental results with high accuracy for all tested ZIPK-FL concentrations (**Fig. 2B**, dashed lines).

To determine to what extent **Model 1** can predict phosphorylation reactions of native myosin by ZIPK, SMM was purified from chicken gizzards, and the phosphorylation reactions were measured. Urea/glycerol PAGE clearly separated 0P-, 1P-, and 2P-MRLCs (**Fig. 6A**), as in the case of the recombinant wild-type MRLC (**Fig. 2B**). Interestingly though, the time courses of the phosphorylation reactions simulated using **Model 1** largely deviated from the experimental results for all tested ZIPK-FL concentrations (**Fig. 6B**, dashed lines). This indicates that **Model 1** (Eqs. (1) and (2); **Fig. 5B**) is not appropriate for SMM phosphorylation and/or the kinetic constants for SMM are deviated from that for MRLC.

**Figure 6.**
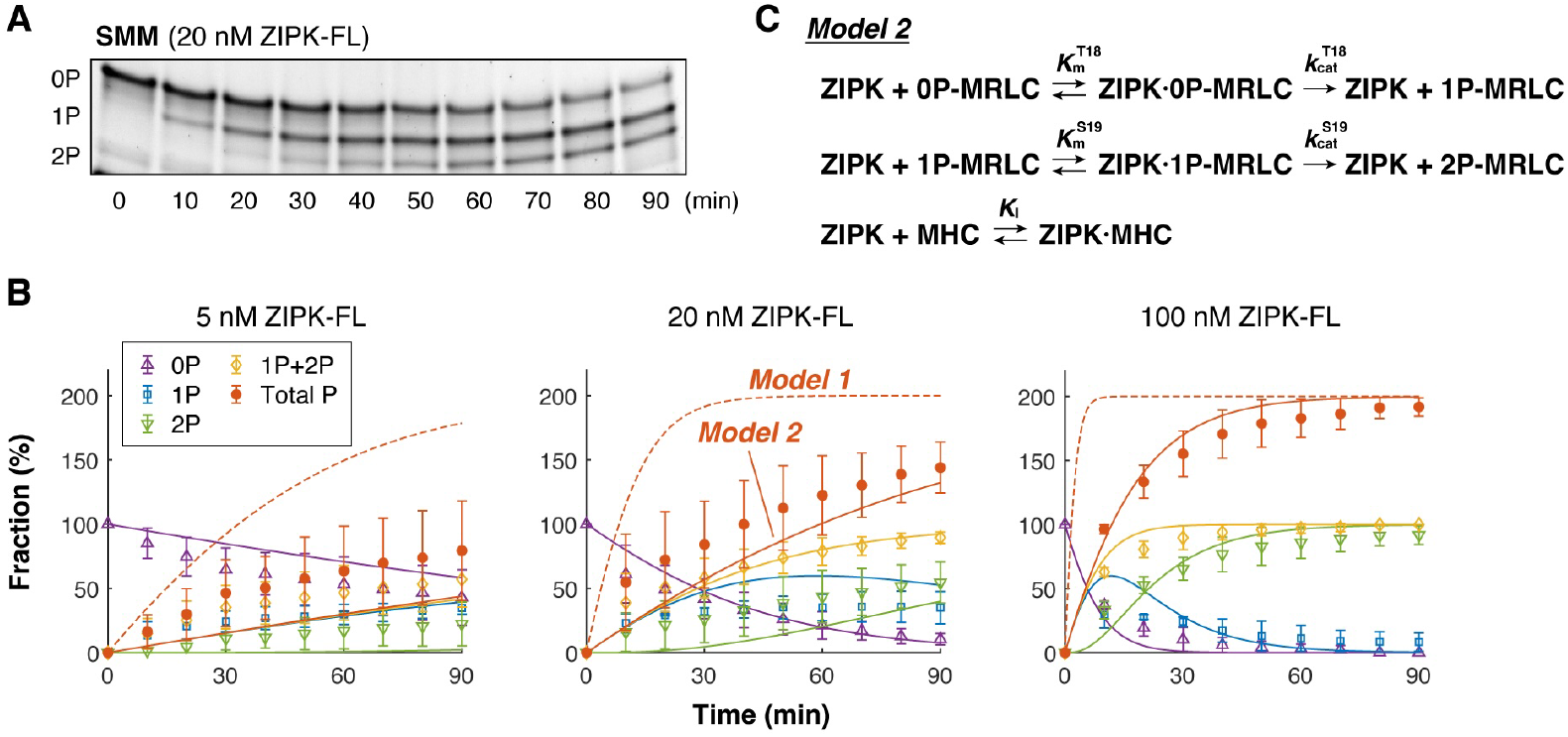
Phosphorylation of SMM by ZIPK-FL. **A**. Images of the urea/glycerol PAGE. One micromolar of smooth muscle myosin (SMM) purified from chicken gizzards was incubated with 20 nM of ZIPK-FL in the presence of 1 mM ATP for the indicated times at 25°C. Subsequently, 0P, 1P, and 2P MRLCs were separated by gel electrophoresis. **B**. Time course of the phosphorylation reactions. Fractions of 0P, 1P, 2P, and 1P+2P MRLCs, determined from densitometry of the urea/glycerol PAGE shown in A, were plotted. Total P indicates the total phosphorylation level, i.e., total P (%) = 1P (%) + 2×2P (%). Three independent experiments were performed. The plots and error bars indicate the mean values and SDs, respectively. Dashed lines represent the time courses predicted from **Model 1** (Eqs. (1) and (2), **Fig. 5B**, right) using the kinetic constants summarized in **Table 1**. Solid lines represent the time courses predicted from **Model 2** shown in C (Eqs. (4) and (5)), using the kinetic constants summarized in **Table 1** and **Table 2. C**. Reaction scheme for the phosphorylation of SMM.

### Measurement of the interactions between ZIPK and SMM

Although the biological species of MRLC and SMM used in the experiments were different (human and chicken, respectively), the amino acid sequence of MRLC is highly conserved between the two species (Identities: 159/172 (92%), BLAST analysis; **Fig. 7A**). In particular, the sequences around Ser^19^ and Thr^18^ are identical (**Fig. 7A**, highlighted in black). Therefore, it is unlikely that the kinetic constants for the phosphorylation reactions largely deviate between human and chicken MRLCs. We confirmed that the phosphorylation rate of chicken MRLC isolated from SMM is comparable to that of recombinant human MRLC, but the phosphorylation rate of chicken MRLC incorporated into SMM was markedly slower (**Fig. 7B**).

**Table 2.** The dissociation constants of ZIPK and SMM. The dissociation constants between ZIPK and SMM were estimated from the co-sedimentation assay.

|  | ZIPK-FL | ZIPK-ΔC |
| --- | --- | --- |
| $K_I$ (μM) | 0.30 | 0.65 |

**Figure 7.**
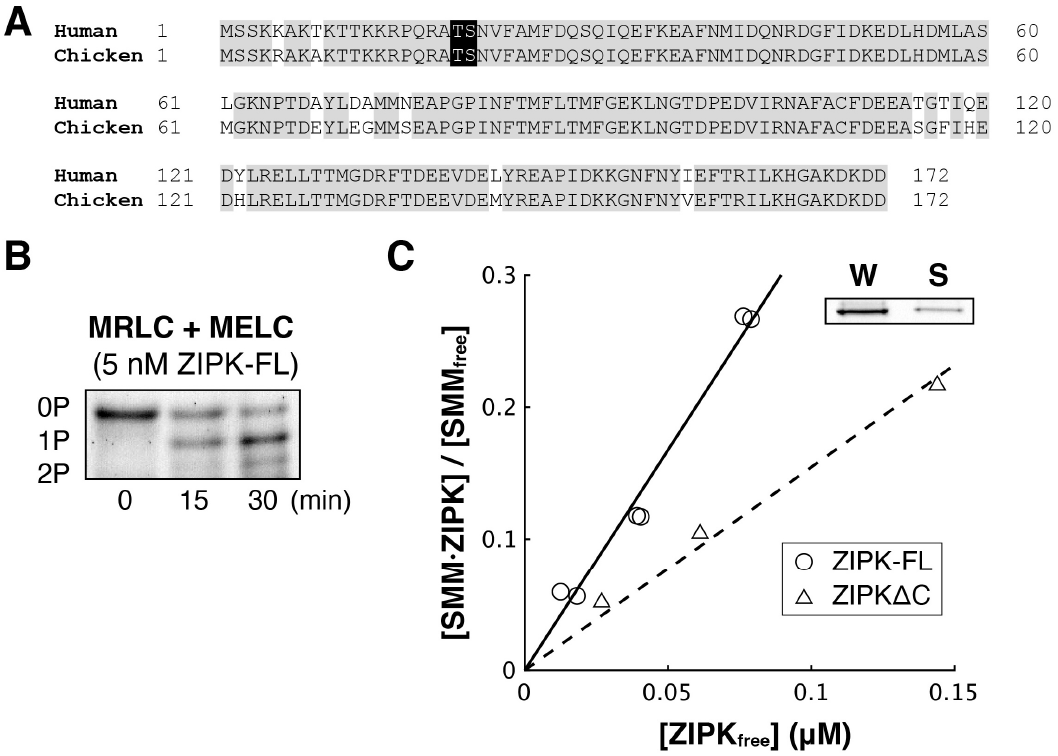
Characterization of the interaction between ZIPK and SMM. **A**. Comparison of the amino acid sequences of MRLC between human and chicken. Gray: identical residues. Black: residues phosphorylated by ZIPK. **B**. Image of urea/glycerol PAGE. A mixture of native MRLC and MELC isolated from chicken gizzard SMM (2 *μ*M) was incubated with 5 nM of ZIPK-FL in the presence of 1 mM ATP for the indicated times at 25°C, then subjected to urea/glycerol PAGE. In total, 50.2%(1P) + 2×30.4%(2P) = 111.0% of MRLC was phosphorylated. This value was comparable to that of wild-type human MRLC (121.3±6.8%; **Fig. 2B**), but largely deviated from that of native chicken gizzard SMM (31.5±4.1%; **Fig. 6B**). **C**. Results of the co-sedimentation experiments of ZIPK and SMM. The solid and dashed lines represent the least-square fitting of the experimental data of ZIPK-FL and ZIPKΔC, respectively. The inverse of the slope corresponds to the dissociation constant *K*I. Inset: A typical example of the image of SDS-PAGE (250 nM ZIPKΔC). The bands corresponding to ZIPK in the uncentrifuged sample (W) and the supernatant (S) are shown.

To account for the difference between the phosphorylation rates of isolated MRLC and MRLC incorporated into SMM, we hypothesized that, in addition to MRLC, ZIPK-FL also interacts with the myosin heavy chain (MHC). Consequently, the competitive interaction with MHC will lower the phosphorylation rate of MRLC. To test this hypothesis, we estimated the affinity of ZIPK-FL for MHC. We assumed that the affinity for MHC was independent of the phosphorylation state of SMM. It is known that SMM oligomerizes to form myosin filaments upon phosphorylation. Therefore, the dissociation constant of ZIPK-FL and SMM can be determined using a co-sedimentation assay. Unphosphorylated SMM (1 *μ*M) was incubated at 25°C for 60 min with ZIPK-FL at various concentrations ranging from 125 nM to 500 nM, to induce phosphorylation and filament formation. Under all conditions, SMM must be almost completely diphosphorylated (2P), as predicted from the experimental results (**Fig. 6B**). After incubation, small aliquots of the samples were taken, then the rests were centrifuged to precipitate myosin filaments. The uncentrifuged samples (W) and the supernatants (S) were subjected to SDS-PAGE (**Fig. 7C**, inset). Upon phosphorylation, > 95% of SMM precipitated under all conditions. The band intensities of ZIPK-FL between W and S were compared to measure the concentrations of bound and unbound ZIPK-FL to SMM ([SMM · ZIPK] and [ZIPK_free_], respectively). Then, [SMM · ZIPK]/[SMM_free_] = [SMM · ZIPK]/([SMM_tot_] − [SMM · ZIPK]) was plotted against the concentration of ZIPK in the supernatant [ZIPK_free_] (**Fig. 7C**). From the least-squares linear fitting of the following equation

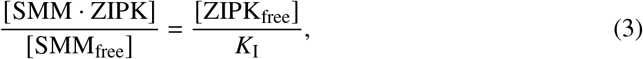

the dissociation constant *K*I of ZIPK-FL and SMM was obtained. The obtained value (*K*I = 0.30 *μ*M; **Table 2**) was nearly one order of magnitude smaller than the Michaelis constant of ZIPK-FL and MRLC (*K*_m_ = 4.2 *μ*M). Although the obtained value includes interactions between ZIPK-FL and MRLC incorporated into SMM, the smaller value suggests that ZIPK-FL preferentially binds to MHC.

### Prediction of the time courses of SMM phosphorylation

The co-sedimentation assay suggests that ZIPK-FL interacts with MHC. Accordingly, it was expected that binding of ZIPK-FL to MHC would reduce the phosphorylation rates of MRLC forming a complex with MHC, compared to isolated MRLC. To account for this effect, we revised **Model 1** by including the competitive binding of ZIPK to MHC (**Fig. 6C**):

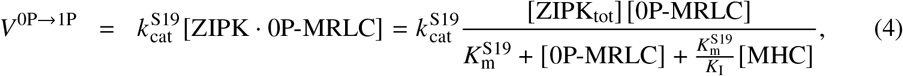

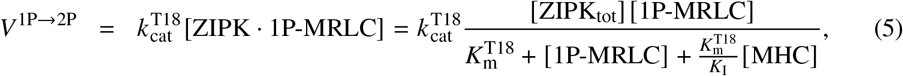

where [MHC] indicates the concentration of MHC. Using the revised model (**Model 2**) and the binding constant *K*_I_ of ZIPK-FL to SMM determined from the co-sedimentation assay (**Table 2**), the time courses of SMM phosphorylation by ZIPK-FL were simulated. The resulting simulated time courses (**Fig. 6B**, solid lines) mostly overlapped with the experimental results, showing that the phosphorylation reactions of SMM by ZIPK-FL can be approximated by **Model 2** (Eqs. (4) and (5); **Fig. 6C**).

### Validation of the kinetic model of SMM phosphorylation

To test the validity of **Model 2**, we prepared a C-terminal deletion mutant of ZIPK (ZIPKΔC) (**Fig. 1**). It has been reported that ZIPK has a short isoform lacking the C-terminus, and this isoform has a higher phosphorylation activity towards SMM[25]. Therefore, it was expected that deletion of the C-terminal domain would affect the phosphorylation kinetics of MRLC and/or the binding affinity of ZIPK for MHC.

First, the kinetic constants of phosphorylation of MRLC at Ser^19^ and Thr^18^ by ZIPKΔC were determined using the same method as that used for ZIPK-FL. We found that both the phosphorylation rates of Ser^19^ and Thr^18^ by ZIPKΔC were higher than those of ZIPK-FL (**Fig. 8**), which is consistent with the case of the short isoform of ZIPK[25]. To maintain consistency with the kinetic model of ZIPK-FL and to keep the model as simple as possible, we assumed that ZIPKΔC phosphorylates Ser^19^ and Thr^18^ with identical Michaelis constants and that the ratio of their catalytic rates is the same as that observed for ZIPK-FL, i.e., 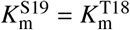 and 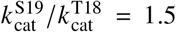. Based on these assumptions, we first determined 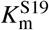 and 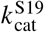 by nonlinear least-squares fitting of the Michaelis–Menten model (Eq.(1)) to the T18A mutant data (**Fig.8**, red plots and curve). Subsequently, the curve for the S19D mutant was simulated under the assumption that 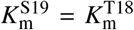 and 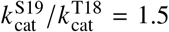. The simulated curve closely matched the experimental data (**Fig. 8**, blue plots and curve), thereby validating the model. The kinetic constants obtained from this analysis (**Table 1**) indicate that both the binding affinity and the catalytic rate of ZIPKΔC are higher than those of ZIPK-FL. Using these kinetic constants and **Model 1**, phosphorylation reactions of wild-type MRLC were simulated and compared with the experimental results (**Fig. 9A**, top, and **Fig. 9B**). We confirmed that **Model 1** accurately predicted the time courses of the phosphorylation reactions of wild-type MRLC by ZIPKΔC (**Fig. 9B**, dashed lines).

**Figure 8.**
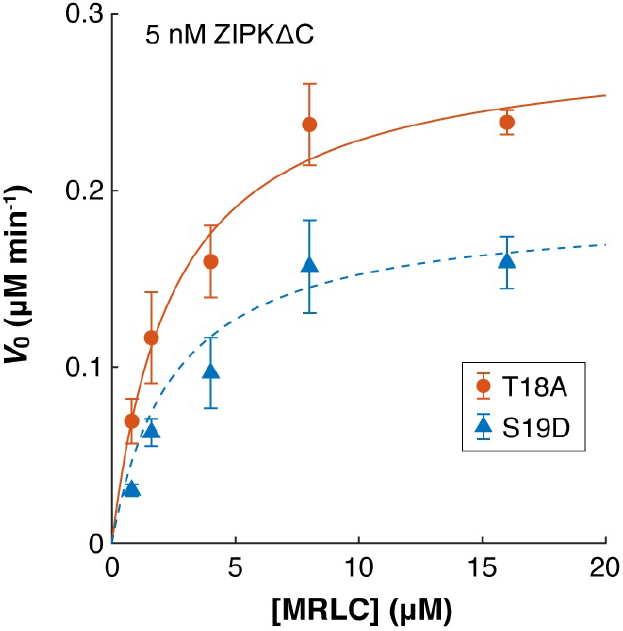
Determination of the kinetic constants for the phosphorylation reactions of MRLC by ZIPKΔC. Phosphorylation rates *V*0 of two MRLC mutants at various MRLC concentrations in the presence of 5 nM ZIPKΔC and 1 mM ATP. The mixtures were incubated at 25°C for 5 min, and the reaction rates were estimated by densitometry of urea/glycerol PAGE. The mean values of the independent experiments were plotted (*n* = 3–8). Error bars indicate SDs. The red solid line indicates the nonlinear fitting of Michaelis–Menten reaction model (Eq. (1)) using 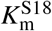 and 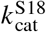 as the fitting parameters. The blue broken line indicates the simulated curve using Michaelis–Menten reaction model (Eq. (2)) and asumptions that 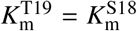 and 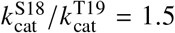. The kinetic constants determined through the nonlinear fittings are summarized in **Table 1**.

**Figure 9.**
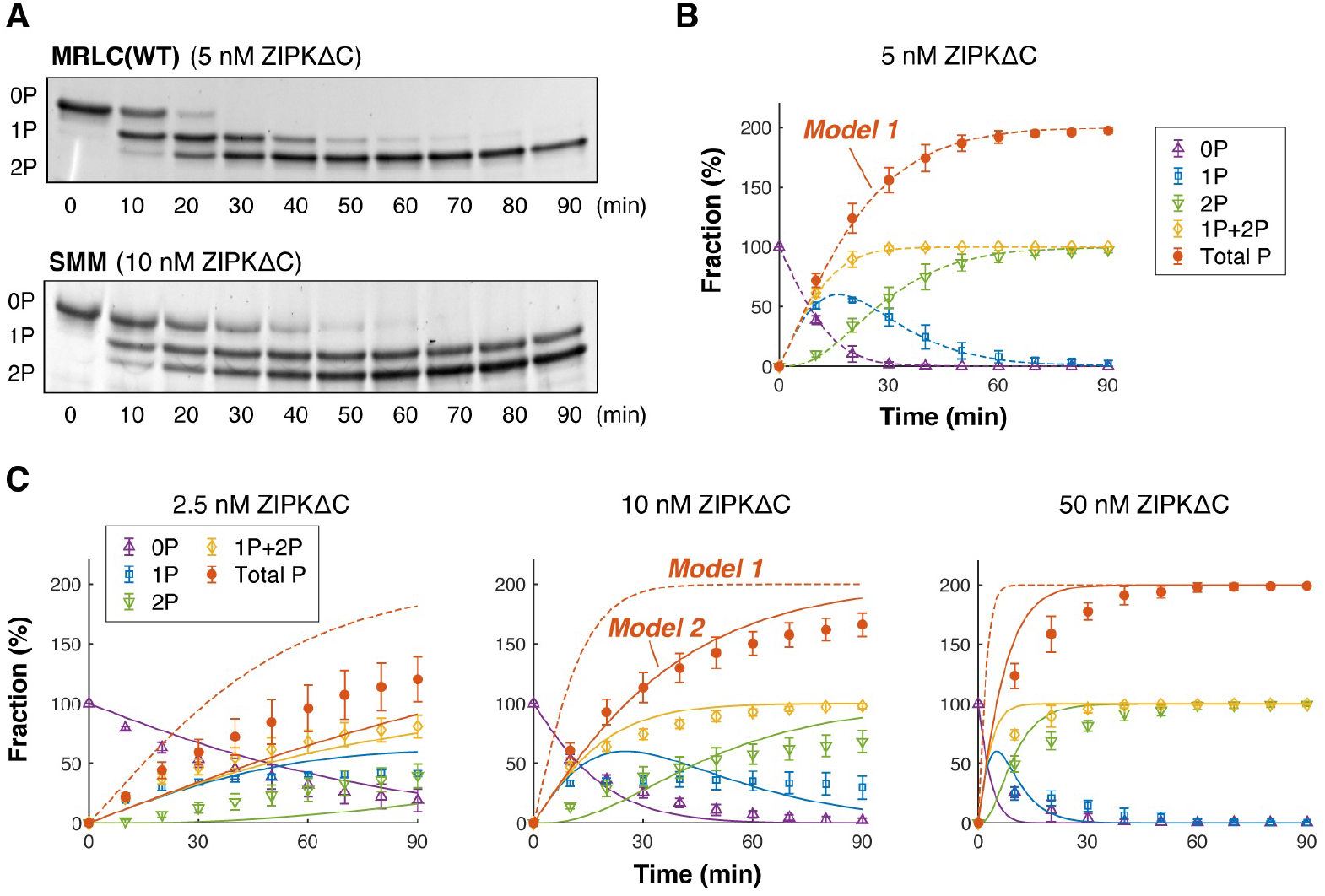
Phosphorylation of wild-type MRLC and SMM by ZIPKΔC. **A**. Images of urea/glycerol PAGE. Wild-type human MRLC (top) or SMM purified from chicken gizzards (bottom) were incubated with 5 nM or 10 nM of ZIPKΔC in the presence of 1 mM ATP for the indicated times at 25°C. The 0P, 1P, and 2P MRLCs were separated by gel electrophoresis. **B, C**. Time courses of the phosphorylation reactions of 2 *μ*M MRLC (**B**) and 1 *μ*M SMM (**C**). Fractions of 0P, 1P, 2P, and 1P+2P MRLCs, determined from densitometry of urea/glycerol PAGE, were plotted. Total P indicates the total phosphorylation level, i.e., total P (%) = 1P (%) + 2×2P (%). Three independent experiments were performed. The plots and error bars indicate the mean values and SDs, respectively. Dashed lines represent the time courses predicted from **Model 1** (Eqs. (1) and (2), **Fig. 5B**, right) using the kinetic constants summarized in **Table 1**. Solid lines represent the time courses predicted from **Model 2** (Eqs. (4) and (5), **Fig. 6C**) using the kinetic constants summarized in **Table 1** and **Table 2**.

Next, we measured the dissociation constant of ZIPKΔC from SMM using a co-sedimentation assay (**Fig. 7C**). The affinity was markedly weakened by the C-terminal deletion (**Table 2**), suggesting that the C-terminal region is involved in binding to MHC. Finally, the phosphorylation reactions of SMM by ZIPKΔC were simulated using **Model 2** (Eqs. (4) and (5), **Fig. 6C**) and compared with the experimental results (**Fig. 9C**). Indeed, **Model 2** accurately predicted the experimental results for all the tested ZIPKΔC concentrations (**Fig. 9C**, solid lines), while the time courses simulated using **Model 1** largely deviated from the experimental results (**Fig. 9C**, dashed lines). This result strengthens the validity of **Model 2**.

## Discussion

ZIPK is phosphorylated at least at six sites, Thr^180^, Thr^225^, Thr^265^, Thr^299^, Thr^306^, and Ser^311^ in the cell to regulate both its enzyme activity and intracellular localization[35, 36]. It is also known that ZIPK undergoes autophosphorylation[15, 35, 25] through activation segment dimerization[37]. Stoichiometric analysis revealed that *∼*6 mol of phosphate/mol of protein is incorporated into ZIPK by autophosphorylation[35], suggesting that six sites are autophosphorylated. Mutational analysis has shown that phosphorylation of Thr^180^, Thr^225^, and Thr^265^ are essential for full autophosphorylation and kinase activity of ZIPK[35, 36]. Pre-incubation of ZIPK with 0.1 mM ATP at 4°C for 5 min resulted in a *∼*3 fold increase in its kinase activity toward the MRLC peptide[35]. The C-terminal truncation mutant at amino acid 273 (ZIPKΔ273) also showed a *∼*3 fold increase by the pre-incubation[35]. In addition, it has been reported that ZIPK has a short isoform lacking the leucine zipper motif, and both the long and short isoforms undergo autophosphorylation[25]. In both isoforms, autophosphorylation was terminated within 1 min at 25°C in the presence of 0.2 mM [*γ*-^32^P]ATP[25]. In the present study, 1 mM ATP was used for all experiments. Therefore, both the full-length ZIPK and the C-terminal truncation mutant were most likely immediately autophosphorylated, and thus the kinetic constants determined in this study can be considered as the constants of fully autophosphorylated ZIPK.

The mechanism of MRLC phosphorylation by ZIPK has been under debate. Hosoya and colleagues first identified ZIPK as an MRLC kinase and proposed a sequential phosphorylation mechanism, in which ZIPK phosphorylates Ser^19^ first, followed by Thr^18^ similar to MLCK, based on two-dimensional phosphopeptide mapping and comparisons with MLCK-phosphorylated MRLC patterns[16]. Later, Ikebe and colleagues proposed a random phosphorylation mechanism: ZIPK phosphorylates Ser^19^ and Thr^18^ in no specific order, supported by good agreement between experimental phosphorylation time courses and predictions from a random phosphorylation model[21]. However, neither study provided direct evidence. In the present study, we directly observed MRLC phosphorylation using mass spectrometry. Quantitative analysis calibrated with synthetic monophosphorylated peptides revealed that ZIPK phosphorylates MRLC first at Ser^19^ and then at Thr^18^, providing direct evidence for a sequential phosphorylation mechanism.

We observed that the phosphorylation rate of SMM was slower than that of isolated MRLC. This result is consistent with previous studies[21, 25], however, the molecular mechanism responsible for this difference remains unresolved. We initially thought that the difference is coming from the folded 10 S conformation of unphosphorylated SMM. SMM undergoes a transformation from the folded 10 S conformation to the extended 6 S conformation upon phosphorylation[38, 39, 40]. The 10 S conformation of SMM is more resistant to proteolysis by papain[41] and trypsin[42] than the 6 S conformation. Therefore, it was expected that the folded 10 S conformation blocked the binding of ZIPK to MRLC. Interestingly though, the co-sedimentation assay indicated that the affinity of ZIPK for the heavy chain might be about one order of magnitude higher than that for the regulatory light chain. This suggests that the observed difference between the phosphorylation rates of isolated MRLC and SMM is primarily due to the competitive binding of ZIPK to the heavy chain of SMM, rather than inhibition of its binding to the regulatory light chain when SMM is folded in the 10 S conformation.

Since ZIPK has a unique domain in its C-terminus among the DAPK family, it is important to understand the role of the C-terminal domain in its regulation. It has been reported that C-terminal truncation at amino acids 273 or 342, or mutations in the leucine zipper motif that inhibit self-association, increased the phosphorylation activity towards the MRLC peptide, suggesting that the C-terminus has a phosphorylation-independent autoinhibitory role[35]. The short isoform of ZIPK lacking the leucine zipper motif has been reported to be a higher kinase activity than the long isoform towards SMM and MRLC[25]. These results are consistent with our results, whereby the C-terminal truncation increases the phosphorylation rates of SMM and MRLC. Furthermore, we found that truncation of the C-terminal domain increases both the affinity for MRLC and the catalytic rates of Ser^19^ and Thr^18^ phosphorylation, while it diminishes the affinity for the heavy chain of SMM. All the observed changes brought on by C-terminal truncation positively contribute to enhanced phosphorylation of SMM, suggesting a role for the C-terminal domain in the regulation of kinase activity.

In addition to MRLC, ZIPK has been shown to interact with ROCK and smooth muscle myosin light chain phosphatase (SMPP-1M). ZIPK forms a complex with ROCK and is phosphorylated at Thr^265^ in HeLa cells during cytokinesis[19], however, the binding domain has not yet been determined. As for SMPP-1M, ZIPK binds to the C-terminal domain of the myosin targeting subunit (MYPT1) of SMPP-1M[43]. Both the long and short isoforms of ZIPK bind to MYPT1, but the mutant consisting of only the kinase domain does not bind to MYPT1, suggesting that residues Ile^278^–Ser^312^, common to the short and long isoforms, are involved in this binding[25]. An *in vitro* study suggests that ZIPK preferentially binds to MYPT1 in the cell because the affinity of ZIPK for MYPT1 is markedly higher than that for ROCK[44]. More precisely, the observed *K*_m_ for MYPT1 is *∼*2 *μ*M, which is approximately 15-fold lower than that determined for ROCK. Here we found that, in addition to MRLC, ROCK, and MYPT1, ZIPK can directly bind to the heavy chain of SMM. Indeed, ZIPK might have a *∼*10 fold and *∼*5 fold higher affinity for SMM than for isolated MRLC and SMPP-1M, respectively, which were estimated from the dissociation constant *K*I for SMM (*∼*0.4 *μ*M; **Table 2**) and the Michaelis constants *K*_m_ for isolated MRLC (*∼*4 *μ*M; **Table 1**) and MYPT1 (*∼*2 *μ*M)[44].

We also found that the affinity of ZIPK for SMM was weakened by truncation of its C-terminal domain. This suggests that the C-terminal domain is involved in the binding of ZIPK to the myosin heavy chain. Consistently, while wild-type ZIPK was localized in the cytosol, the C-terminal truncation mutants (ZIPKΔ273 and ZIPKΔ342) showed diffuse staining in HeLa cells[35]. The short isoform of ZIPK showed a similar tendency in ARPE19 cells[25]. Conversely, using GST-pull down assays and C-terminal deletion mutants, it was shown that the kinase domain can directly bind to SMM[25]. However, it is not known whether the kinase domain binds to the heavy chain of SMM. Further investigation is needed to determine the binding domain(s) of ZIPK to the heavy chain of SMM and the interacting domain(s) in the heavy chain. Notably, it is important to determine whether the binding site of ZIPK on the myosin heavy chain is shared with MYPT1 and thus the binding of ZIPK competes with MYPT1. If the binding of MYPT1 and ZIPK to the myosin heavy chain competes with each other, this competition might facilitate efficient on/off switching of myosin contractility in the cell by controlling localizations of the kinase (ZIPK) and the phosphatase (SMPP-1M) in a mutually exclusive manner.

## Conclusion

In the present study, we determined the reaction scheme of the phosphorylation of SMM by ZIPK. First, we found that ZIPK phosphorylates MRLC sequentially, first at Ser^19^ and then at Thr^18^, as determined by quantitative mass spectrometry of wild-type MRLC. Second, using the phosphomimic and unphosphorylatable mutants, we estimated by electrophoresis that the phosphorylation rate at Ser^19^ on unphosphorylated MRLC is 1.5 times faster than that at Thr^18^ on Ser^19^-phosphorylated MRLC. Third, we observed that the phosphorylation rate of SMM was slower than that of isolated MRLC. Then, we proposed that competitive binding of ZIPK to MRLC and the myosin heavy chain results in suppression of the phosphorylation reaction of SMM. Finally, we demonstrated that both the time courses of the phosphorylation reactions of SMM and isolated MRLC were reproduced by a simple kinetic model with high accuracy using the kinetic constants determined from independent experiments, validating the proposed kinetic models. Taken together, the present study successfully determines the reaction schemes of the phosphorylation of isolated MRLC and SMM by ZIPK. This will contribute to understanding the regulatory mechanism of myosin contractility in the cell, which is organized by the cooperative action of myosin kinases and phosphatases.

## Author Contributions

M.M. designed the research. M.Y., L.T.T., and M.M. performed phosphorylation assays and co-sedimentation assays. R.N. and Y.S. performed mass spectrometry analysis. M.M. developed kinetic models. All the authors discussed the results. M.M. wrote the manuscript.

## Acknowledgements

Plasmids encoding full-length human ZIPK (pGEX-5X-hZIPK) and wild-type human MRLC2 (pGEX-6P-hMRLC2) are kind gifts from Kozue Hamao (Hiroshima University, Japan). We thank Tomoko Yamanaka (Kyoto University), Hibiki Sakata (Kyoto University), Takaya Izumi (Kyoto University), and Kanako Takamura (RIKEN) for their help with the experiments. This work was partly supported by JST PRESTO, Japan (grant no. JPMJPR20ED to M.M.), Grantin-Aid for Transformative Research Areas (A) (grant no. 22H05171 to M.M.) and Grant-in-Aid for Challenging Research (Exploratory) (grant no. 21K19220 to M.M.) from the Ministry of Education, Culture, Sports, Science, and Technology, Japan, Kishimoto Foundation Research Grant (to M.M.), Takeda Science Foundation (to M.M.), SPIRITS 2020 of Kyoto University (to M.M.), and The Hakubi project of Kyoto University (to M.M.).

---

**Supplementary Figure S1.**
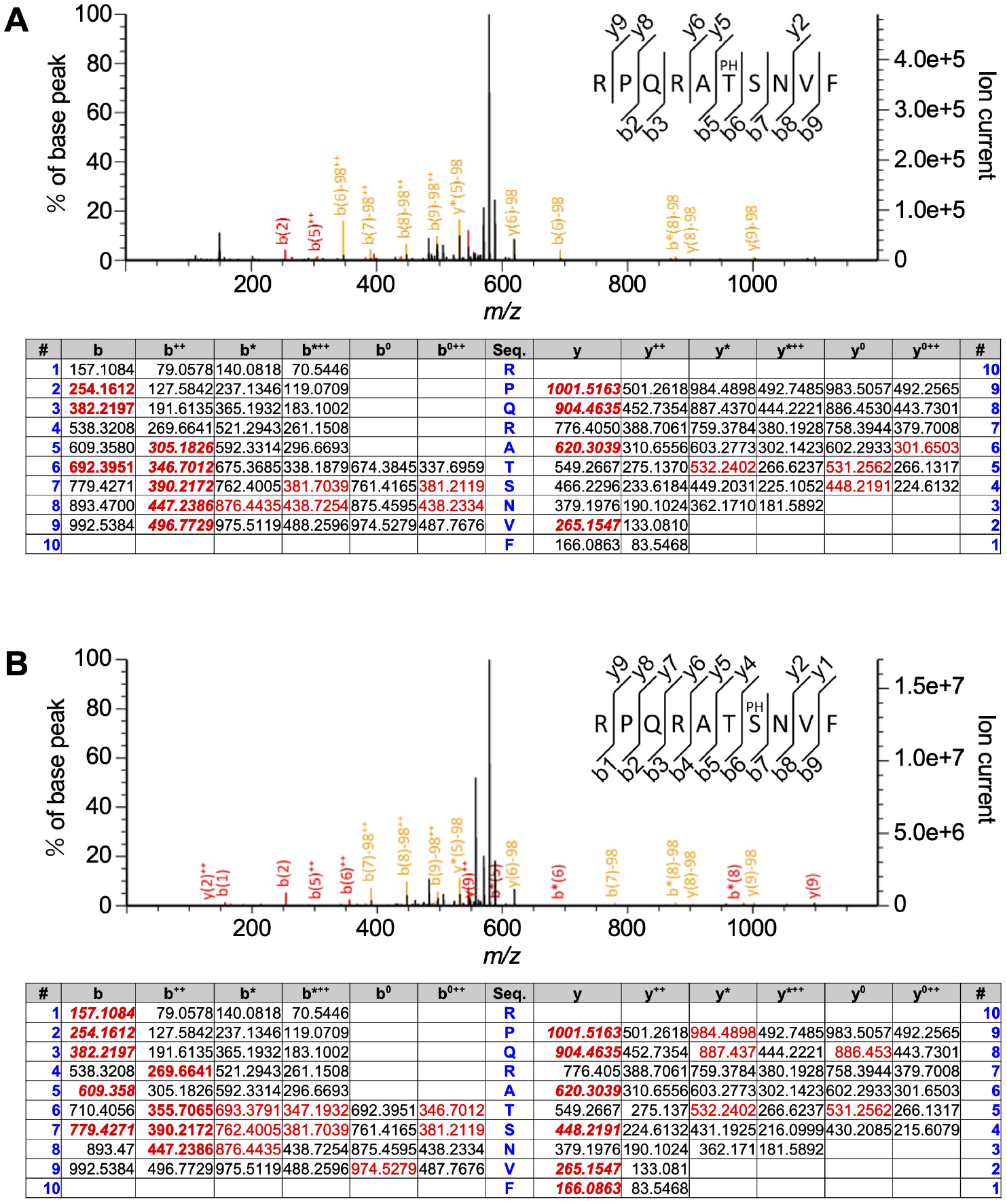
MS/MS analysis of ZIPK-FL dependent phosphorylation of wild-type MRLC. MS/MS spectra of the peptide RPQRATSNVF, monophosphorylated at Thr^18^ (**A**) and Ser^19^ (**B**), along with the summary table of b- and y-type fragment ions are shown. Retention times: 18.4 min (**A**) and 19.4 min (**B**). Red: Ions matched to the spectrum. Bold red: Scoring ion series with more matches than expected by chance. Bold italic red: Significant ion series contributing to the ion score. Red peaks in the spectra indicate peaks matching the summary table; yellow peaks indicate peaks generated by the neutral loss of a phosphate group.

## References

[1] Howard, J. (2001) Mechanics of motor proteins and the cytoskeleton. Sinauer Associates, Inc., Sunderland, MA

[2] Vicente-Manzanares, M., Ma, X., Adelstein, R. S., and Horwitz, A. R. (2009) Non-muscle myosin II takes centre stage in cell adhesion and migration. Nat. Rev. Mol. Cell Biol. 10, 778–790

[3] Somlyo, A. P. and Somlyo, A. V. (1994) Signal transduction and regulation in smooth muscle. Nature 372, 231–236

[4] Ikebe, M., Hartshorne, D. J., and Elzinga, M. (1986) Identification, phosphorylation, and dephosphorylation of a second site for myosin light chain kinase on the 20,000-dalton light chain of smooth muscle myosin. J. Biol. Chem. 261, 36–39

[5] Ikebe, M., Koretz, J., and Hartshorne, D. J. (1988) Effects of phosphorylation of light chain residues threonine 18 and serine 19 on the properties and conformation of smooth muscle myosin. J. Biol. Chem. 263, 6432–6437

[6] Ikebe, M. and Hartshorne, D. J. (1985) Phosphorylation of smooth muscle myosin at two distinct sites by myosin light chain kinase. J. Biol. Chem. 260, 10027–10031

[7] Sellers, J. R., Spudich, J. A., and Sheetz, M. P. (1985) Light chain phosphorylation regulates the movement of smooth muscle myosin on actin filaments. J. Cell Biol. 101, 1897–1902

[8] Kamisoyama, H., Araki, Y., and Ikebe, M. (1994) Mutagenesis of the phosphorylation site (Serine 19) of smooth muscle myosin regulatory light chain and its effects on the properties of myosin. Biochemistry 33, 840–847

[9] Umemoto, S., Bengur, A. R., and Sellers, J. R. (1989) Effect of multiple phosphorylations of smooth muscle and cytoplasmic myosins on movement in an in vitro motility assay. J. Biol. Chem. 264, 1431–1436

[10] Hong, F., Haldeman, B. D., Jackson, D., Carter, M., Baker, J. E., and Cremo, C. R. (2011) Biochemistry of smooth muscle myosin light chain kinase. Arch. Biochem. Biophys. 510, 135–146

[11] Hathaway, D. R. and Adelstein, R. S. (1979) Human platelet myosin light chain kinase requires the calcium-binding protein calmodulin for activity. Proc. Natl. Acad. Sci. U. S. A. 76, 1653–1657

[12] Adelstein, R. S. and Klee, C. B. (1981) Purification and characterization of smooth muscle myosin light chain kinase. J. Biol. Chem. 256, 7501–7509

[13] Amano, M., Ito, M., Kimura, K., Fukata, Y., Chihara, K., Nakano, T., Matsuura, Y., and Kaibuchi, K. (1996) Phosphorylation and activation of myosin by Rho-associated kinase (Rho-kinase). J. Biol. Chem. 271, 20246–20249

[14] Amano, M., Nakayama, M., and Kaibuchi, K. (2010) Rho-kinase/ROCK: A key regulator of the cytoskeleton and cell polarity. Cytoskeleton 67, 545–554

[15] Kawai, T., Matsumoto, M., Takeda, K., Sanjo, H., and Akira, S. (1998) ZIP Kinase, a novel serine/threonine kinase which mediates apoptosis. Mol. Cell. Biol. 18, 1642–1651

[16] Murata-Hori, M., Suizu, F., Iwasaki, T., Kikuchi, A., and Hosoya, H. (1999) ZIP kinase identified as a novel myosin regulatory light chain kinase in HeLa cells. FEBS Lett. 451, 81–84

[17] Komatsu, S. and Ikebe, M. (2004) ZIP kinase is responsible for the phosphorylation of myosin II and necessary for cell motility in mammalian fibroblasts. J. Cell Biol. 165, 243–254

[18] Hosoba, K., Komatsu, S., Ikebe, M., Kotani, M., Wenqin, X., Tachibana, T., Hosoya, H., and Hamao, K. (2015) Phosphorylation of myosin II regulatory light chain by ZIP kinase is responsible for cleavage furrow ingression during cell division in mammalian cultured cells. Biochem. Biophys. Res. Commun. 459, 686–691

[19] Hamao, K., Ono, T., Matsushita, M., and Hosoya, H. (2020) ZIP kinase phosphorylated and activated by Rho kinase/ROCK contributes to cytokinesis in mammalian cultured cells. Exp. Cell. Res. 386, 111707

[20] Murata-Hori, M., Fukuta, Y., Ueda, K., Iwasaki, T., and Hosoya, H. (2001) HeLa ZIP kinase induces diphosphorylation of myosin II regulatory light chain and reorganization of actin filaments in nonmuscle cells. Oncogene 20, 8175–8183

[21] Niiro, N. and Ikebe, M. (2001) Zipper-interacting protein kinase induces Ca^2+^-free smooth muscle contraction via myosin light chain phosphorylation. J. Biol. Chem. 276, 29567–29574

[22] Moffat, L. D., Brown, S. B. A., Grassie, M. E., Ulke-Lemeé, A., Williamson, L. M., Walsh, M. P., and MacDonald, J. A. (2011) Chemical genetics of zipper-interacting protein kinase reveal myosin light chain as a bona fide substrate in permeabilized arterial smooth muscle. J. Biol. Chem. 286, 36978–36991

[23] Shohat, G., Shani, G., Eisenstein, M., and Kimchi, A. (2002) The DAP-kinase family of proteins: study of a novel group of calcium-regulated death-promoting kinases. Biochim. Biophys. Acta. 1600, 45–50

[24] Shiloh, R., Bialik, S., and Kimchi, A. (2014) The DAPK family: a structure-function analysis. Apoptosis 19, 286–297

[25] Takamoto, N., Komatsu, S., Komaba, S., Niiro, N., and Ikebe, M. (2006) Novel ZIP kinase isoform lacks leucine zipper. Arch. Biochem. Biophys. 456, 194–203

[26] Perrie, W. T. and Perry, S. V. (1970) An electrophoretic study of the low-molecular-weight components of myosin. Biochem. J. 119, 31–38

[27] Kondo, T., Isoda, R., Uchimura, T., Sugiyama, M., Hamao, K., and Hosoya, H. (2012) Diphosphorylated but not monophosphorylated myosin II regulatory light chain localizes to the midzone without its heavy chain during cytokinesis. Biochem. Biophys. Res. Commun. 417, 686–691

[28] Yamamoto, K. and Miyazaki, M. (2025) Optogenetic actin network assembly on lipid bilayer uncovers the network density-dependent functions of actin-binding proteins. Nat. Commun. in press

[29] Ikebe, M. and Hartshorne, D. J. (1985) Effects of Ca^2+^on the conformation and enzymatic activity of smooth muscle myosin. J. Biol. Chem. 260, 13146–13153

[30] Hughes, C. S., Moggridge, S., Müller, T., Sorensen, P. H., Morin, G. B., and Krijgsveld, J. (2019) Single-pot, solid-phase-enhanced sample preparation for proteomics experiments. Nat. Protoc. 14, 68–85

[31] MacLean, B., Tomazela, D. M., Shulman, N., Chambers, M., Finney, G. L., Frewen, B., Kern, R., Tabb, D. L., Liebler, D. C., and MacCoss, M. J. (2010) Skyline: an open source document editor for creating and analyzing targeted proteomics experiments. Bioinformatics. 26, 966–968

[32] Good, M. C., Vahey, M. D., Skandarajah, A., Fletcher, D. A., and Heald, R. (2013) Cytoplasmic volume modulates spindle size during embryogenesis. Science 342, 856–860

[33] Miyazaki, M., Chiba, M., Eguchi, H., Ohki, T., and Ishiwata, S. (2015) Cell-sized spherical confinement induces the spontaneous formation of contractile actomyosin rings in vitro. Nat. Cell Biol. 17, 480–489

[34] Sakamoto, R., Tanabe, M., Hiraiwa, T., Suzuki, K., Ishiwata, S., Maeda, Y. T., and Miyazaki, M. (2020) Tug-of-war between actomyosin-driven antagonistic forces determines the positioning symmetry in cell-sized confinement. Nat. Commun. 11, 3063

[35] Graves, P. R., Winkfield, K. M., and Haystead, T. A. J. (2005) Regulation of zipper-interacting protein kinase activity in vitro and in vivo by multisite phosphorylation. J. Biol. Chem. 280, 9363–9374

[36] Haystead, T. A. J. (2005) ZIP kinase, a key regulator of myosin protein phosphatase 1. Cell. Signal. 17, 1313–1322

[37] Pike, A. C. W., Rellos, P., Niesen, F. H., Turnbull, A., Oliver, A. W., Parker, S. A., Turk, B. E., Pearl, L. H., and Knapp, S. (2008) Activation segment dimerization: a mechanism for kinase autophosphorylation of non-consensus sites. EMBO J. 27, 704–714

[38] Onishi, H., Suzuki, H., Nakamura, K., Takahashi, K., and Watanabe, S. (1978) Adenosine triphosphatase activity and “thick filament” formation of chicken gizzard myosin in low salt media. J. Biochem. 83, 835–847

[39] Suzuki, H., Onishi, H., Takahashi, K., and Watanabe, S. (1978) Structure and function of chicken gizzard myosin. J. Biochem. 84, 1529–1542

[40] Trybus, K. M., Huiatt, T. W., and Lowey, S. (1982) A bent monomeric conformation of myosin from smooth muscle. Proc. Natl. Acad. Sci. U. S. A. 79, 6151–6155

[41] Onishi, H. and Watanabe, S. (1984) Correlation between the papain digestibility and the conformation of 10S-myosin from chicken gizzard. J. Biochem. 95, 899–902

[42] Ikebe, M. and Hartshorne, D. J. (1984) Conformation-dependent proteolysis of smooth-muscle myosin. J. Biol. Chem. 259, 11639–11642

[43] Endo, A., Surks, H. K., Mochizuki, S., Mochizuki, N., and Mendelsohn, M. E. (2004) Identification and characterization of zipper-interacting protein kinase as the unique vascular smooth muscle myosin phosphatase-associated kinase. J. Biol. Chem. 279, 42055–42061

[44] MacDonald, J. A., Borman, M. A., Murányi, A., Somlyo, A. V., Hartshorne, D. J., and Haystead, T. A. J. (2001) Identification of the endogenous smooth muscle myosin phosphatase-associated kinase. Proc. Natl. Acad. Sci. U. S. A. 98, 2419–2424

